# Macrophages protect against loss of adipose tissue during cancer cachexia

**DOI:** 10.1101/427963

**Authors:** Merve Erdem, Diana Möckel, Sandra Jumpertz, Cathleen John, Athanassios Fragoulis, Ines Rudolph, Johanna Wulfmeier, Jochen Springer, Henrike Horn, Marco Koch, Georg Lurje, Twan Lammers, Gregory van der Kroft, Felix Gremse, Thorsten Cramer

## Abstract

Cancer cachexia represents a central obstacle in medical oncology as it is associated with poor therapy response and reduced overall survival. Systemic inflammation is considered to be a key driver of cancer cachexia, however, clinical studies with anti-inflammatory drugs failed to show a robust cachexia-inhibiting effect. To address this contradiction, we investigated the functional importance of innate immune cells for hepatocellular carcinoma (HCC)-associated cachexia. To this end, we used a transgenic HCC mouse model intercrossed with mice harboring a defect in myeloid cell-mediated inflammation. We identified robust cachexia in the HCC mouse model as evidenced by a marked loss of visceral fat and lean mass. Computed tomography-based analyses demonstrated that a subgroup of human HCC patients displays reduced visceral fat mass, complementing the murine data. While the myeloid cell-mediated inflammation defect resulted in reduced expression of pro-inflammatory cytokines in the serum of HCC-bearing mice, this unexpectedly did not translate into diminished, but rather enhanced cachexia-associated fat loss. Defective myeloid cell-mediated inflammation was associated with decreased macrophage abundance in visceral adipose tissue, suggesting a role for local macrophages in the regulation of cancer-induced fat loss. Taken together, myeloid cell-mediated inflammation displays a rather unexpected beneficial function in a murine HCC model. These results demonstrate that immune cells are capable of protecting the host against cancer-induced tissue wasting, adding a further layer of complexity to the pathogenesis of cachexia and providing a potential explanation for the contradictory results of clinical studies with anti-inflammatory drugs.

## Introduction

Cachexia is a multi-factorial metabolic syndrome characterized by weight loss due to depletion of muscle with or without loss of fat (1). Systemic inflammation, insulin resistance, enhanced muscle protein breakdown and sympathetic nervous system activation are hallmarks of cachexia. A plethora of diseases are associated with cachexia, e.g. chronic infections (HIV, tuberculosis), chronic heart failure, chronic obstructive lung disease and chronic kidney failure (2). The most common association, however, exists between cachexia and cancer, where it can occur in up to 80% of cases (3). Cancer-associated cachexia (CAC) goes along with unfavorable prognosis and plays a causal role in up to 20% of cancer-related deaths (3). CAC is unresponsive to nutritional support and while a lot of progress has been made in the past years regarding the mechanisms of CAC, an effective treatment option is still missing.

Patients suffering from hepatocellular carcinoma (HCC) typically show loss of muscle mass and strength (termed sarcopenia), resulting in frailty and debilitating physical weakness (4). In addition, cachexia is a common characteristic of HCC patients which not only reduces the quality of life, but -together with sarcopenia-impacts significantly on overall prognosis (5) and clinical decision making: Mortality of HCC patients after intra-arterial therapy (6) and liver transplantation (7) as well as dose-limiting toxicities of sorafenib (8), which is the gold standard oral treatment for non-resectable HCC, are independently predicted by sarcopenia. Taken together, cachexia and sarcopenia are of pivotal importance for both patients (quality of life, prognosis) and clinicians (progression, treatment decision).

Against this background, a better understanding of the molecular and cell biological mechanisms that govern HCC-associated sarcopenia and cachexia is urgently needed. As cachexia is a multi-factorial syndrome affecting various organs and cellular systems, this can only be achieved by using *in vivo* model systems that recapitulate the syndrome as a whole (9). With respect to HCC-associated cachexia, the most widely applied system is the rat ascites hepatoma Yoshida AH-130 model. This is characterized by a hypercatabolic state and marked depletion of both skeletal muscle and adipose tissue (10, 11). While the Yoshida AH-130 model is certainly of great value, especially for the identification of potential cachexia-inhibiting drugs (12, 13), a better understanding of the pathogenesis of HCC-associated cachexia is limited by the absence of a practicable murine model system. Intercrossing such a mouse model with conventional or cell type-specific knock-out mice would enable researchers to address a number of hypotheses and would undoubtedly result in a much better understanding of the molecular and cell biological mechanisms that govern HCC-associated cachexia (14).

Here, we describe a robust cachexia phenotype in a transgenic murine HCC model. Intercrosses with mice harboring defective myeloid cell-mediated inflammation unexpectedly resulted in enhanced cachexia-associated loss of adipose tissue even though systemic levels of pro-inflammatory cytokines were lower in the knock-out mice. Furthermore, we present experimental data arguing for a protective role of macrophages in the context of CAC-associated fat loss. Taken together, our results challenge the general understanding of pro-inflammatory cytokines as causal agents of CAC and establish a functional importance of macrophages in the setting of CAC-associated fat loss that has not been previously appreciated.

## Results

### ASV-B mice display robust cancer-associated cachexia

The transgenic ASV-B mouse line is a well-established HCC model based on hepatocyte-specific expression of the SV40 large T oncogene (15). In this model, mice develop dysplastic hepatocytes at 8, hepatic adenomas at 12 and hepatocellular carcinomas at 16 weeks of age (16). We initially evaluated the HCC progression by measuring liver weight in different age groups. As shown in figure 1A, liver weight of ASV-B mice strongly increased with age compared to tumor-free C57BL/6J controls, reflecting tumor progression. Rather unexpectedly, this pronounced increase of liver mass did not affect total body weight. In fact, ASV-B mice were even slightly lighter than tumor-free C57BL/6J mice between 12 and 18 weeks of age (Fig. 1B). Food and water intake were not different between control and HCC-bearing mice, ruling out a functional relevance of anorexia in this setting (Fig. 1C). Altogether, the observed phenotype led us to consider tissue wasting and cachexia in the course of ASV-B tumor formation. To address this, we initially evaluated the mass of different skeletal muscle regions. As can be seen in figure 1D, tumor-bearing mice showed a significant decrease in mass of gastrocnemius (GC), tibialis anterior (TA) and extensor digitorum longus (EDL) muscle over time. Furthermore, heart weight was diminished in ASV-B compared to tumor-free mice in all age groups (Fig. 1D). Along with the muscle wasting, we observed a striking loss of adipose tissue in tumor-bearing mice. Measurement of gonadal fat depots revealed that fat loss started at 12 weeks of age and continued throughout tumor progression (Fig. 1E). A representative image of an ASV-B mouse at the age of 16 weeks shows visibly smaller gonadal fat depots and clear tumor nodules in the massively enlarged liver (Fig. 1F). The loss of adipose tissue and muscle mass illustrates the development of cancer-associated cachexia (CAC) in ASV-B mice.

**Figure 1.**
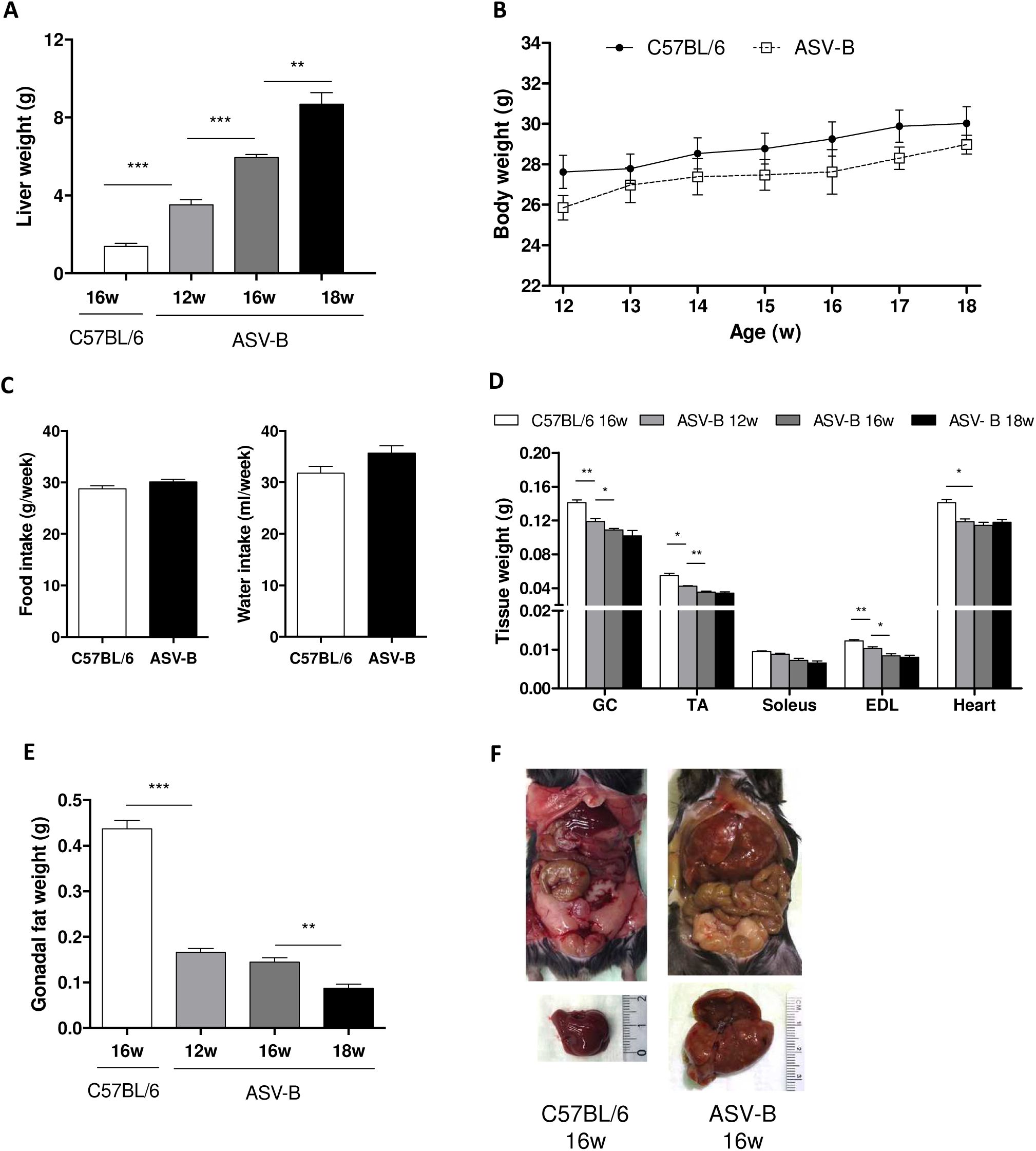
Characterization of cachexia in ASV-B mice. **A**, liver weight and **B**, total body weight of C57BL/6 and ASV-B mice *(n* = 8 per group) were measured at the indicated time points. **C**, food and water intake were measured weekly from 8 to 18 weeks of age in control (n = 3) and ASV-B (n = 8) mice. **D**, different muscle parts (GC: Gastrocnemius, TA: Tibialis anterior, EDL: Extensor digitorum longus) were dissected and weighed for C57BL/6 mice (n = 3) and ASV-B mice at the indicated time points (n = 8, 5, 4 in 12, 16, and 18 weeks, respectively). **E**, the gonadal fat pad was removed and measured at *(n* = 8 per group) at the same time points used in **D**. Data represent means with s.e.m. *P < 0.05; **P < 0.01; *** *P* < 0.001 according to two-way ANOVA (**A**), one-way ANOVA followed by Tukey post hoc test (**B**, **D, E**) or unpaired Student’s T-test (**C**). **F** shows a general view of the abdominal cavity (above) and the resected livers (below) of control and ASV-B mice at the 16 week time point.

### Defective myeloid cell-mediated inflammation unexpectedly aggravates loss of adipose tissue in HCC-bearing mice

Systemic inflammation is widely appreciated as a driving force of cachexia (17). Myeloid cells, e.g. granulocytes, monocytes and macrophages, are the chief cellular effectors of the innate immune system and centrally involved in cancer-associated inflammation (18). Earlier work by us and others has identified the hypoxia-inducible transcription factor HIF-1 as an essential regulator of myeloid cell-mediated inflammation (19). We sought to investigate the functional importance of myeloid cells for the pathogenesis of CAC in ASV-B mice. To this end, we intercrossed ASV-B mice with myeloid cell-specific *Hif1a* knock-out mice (termed ASV-B/Hif1a^MC^) and analyzed body weight and body composition. As can be seen in figure 2A, ASV-B/Hif1a^MC^ mice displayed a non-significant tendency for higher body weight than WT mice. Body composition analysis by nuclear magnetic resonance (NMR) spectroscopy showed a lower amount of total body fat in ASV-B/Hif1a^MC^ mice, again without reaching statistical significance (Fig. 2B). In addition to NMR analyses, *in vivo* micro computed tomography (μCT) imaging of mice was performed to visualize and quantify body composition. 2D cross-sectional images and three-dimensional volume renderings of segmented bones, lungs, liver and fat were obtained. In figure 2C, representative μCT images display fat loss in all depots as well as liver enlargement in tumor-bearing mice. Quantification of the volume analysis indicated a significant decrease of fat amount in ASV-B mice compared to controls (Fig. 2D). The total fat amount in ASV-B/Hif1a^MC^ mice tended to be lower than in ASV-B WT mice (significance level of 0.05, Fig. 2D), which is consistent with the obtained NMR body composition results. Of note, skeletal muscle and heart weight did not differ between ASV-B WT and ASV-B/Hif1a^MC^ mice (Suppl. Fig. 1). Taken together, the defective myeloid cell-mediated inflammation did not result in reduced CAC, but unexpectedly aggravated the CAC-associated fat loss.

**Figure 2.**
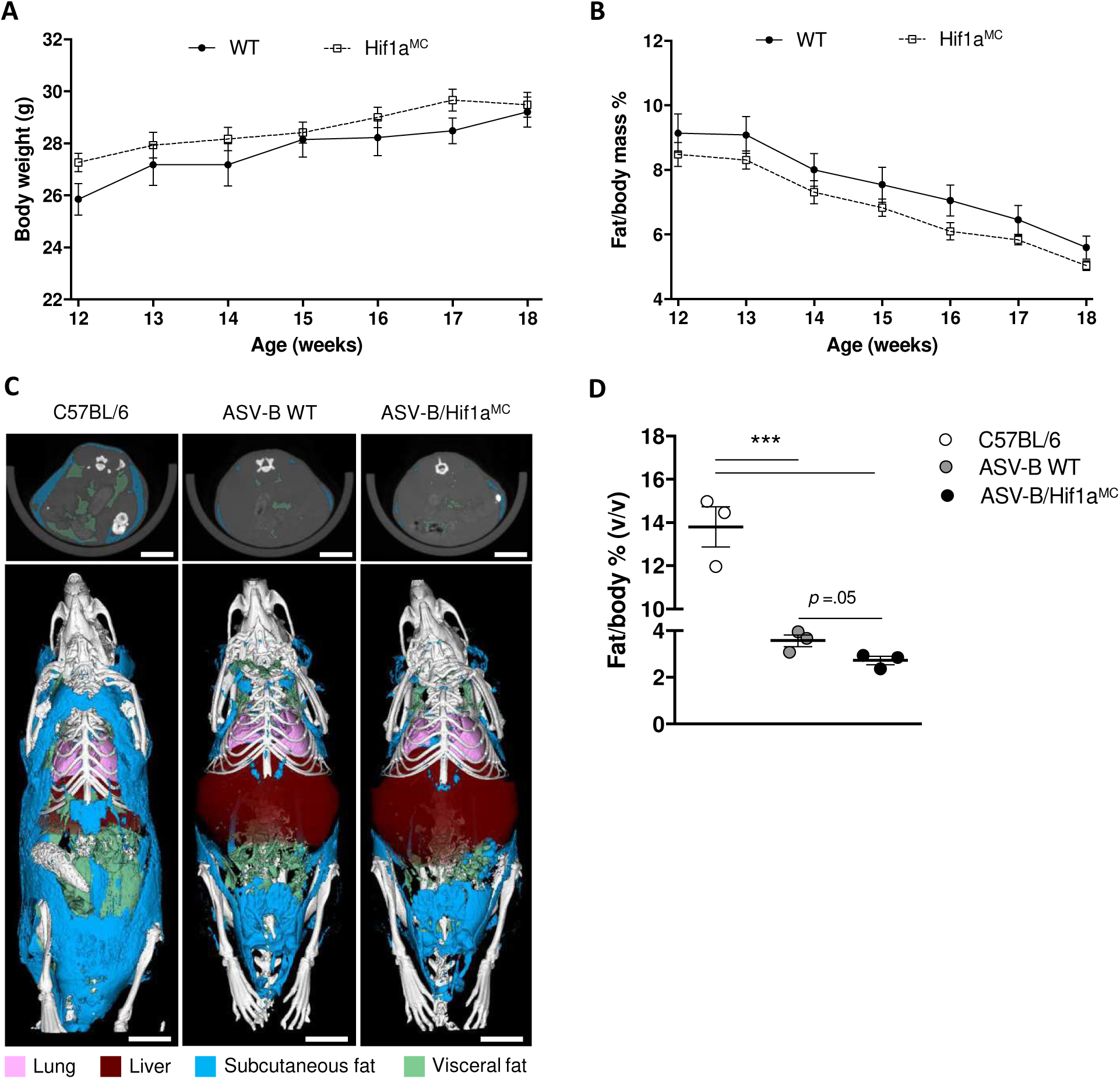
Myeloid cell-specific *Hif1a* KO mice show aggravated fat loss in ASV-B mice. **A**, Body weight of ASV-B WT *(n* = 8) and Hif1a^MC^ *(n* = 6) mice over time. **B**, weekly follow-up NMR analysis (n = 7 per group) for body fat quantification. **C**, μCT imaging at 14 weeks old C57BL/6 and ASV-B WT and Hif1a^MC^ mice; upper panel shows representative 2D cross-sectional μCT images in transversal planes of the abdomen of one representative mouse from each group (subcutaneous and visceral fat tissue is indicated in blue and green, respectively). Lower panel, representative images of three-dimensional volume renderings of segmented bones (white), lungs (pink), liver (brown) and fat (blue/green) upon *in vivo* μCT imaging, scale bar 1 cm. **D**, Quantification of fat volume via μCT imaging (n = 3 per group). Data show means with s.e.m. *\*P* < 0.05; **P < 0.01; *** *P* < 0.001 according to two-way ANOVA (**A, B**) and one-way ANOVA followed by Tukey post hoc test (**D**).

### Visceral adipose tissue of ASV-B mice displays typical cachexia-associated changes

We sought to identify the mechanisms underlying the loss of visceral adipose tissue (VAT) in ASV-B mice and addressed the hypothesis that enhanced lipid mobilization takes place in HCC-bearing mice (20, 21). To this end, we quantified the cell size of adipocytes in epididymal white adipose tissue (eWAT) and could show substantial cell shrinking between control and ASV-B mice (Fig. 3A). ASV-B/Hif1a^MC^ mice had a higher frequency of smaller adipocytes (<1.500 μm^2^) and lower frequency of relatively larger adipocytes (1.500-4.000 μm^2^) than ASV-B WT mice (statistically significant at 2.000 μm^2^) (Suppl. Fig. 2). It recently became clear that white adipose tissue (WAT) is able to switch to a thermogenic fat burning phenotype (termed “browning”) (22). This process was found to contribute to the increased energy expenditure typical for cachexia in different mouse models of cancer cachexia (22, 23). We found significantly elevated mRNA levels of various browning marker genes in WAT of ASV-B mice in comparison to tumor-free controls (Fig. 3B), demonstrating WAT browning in this HCC model. Of note, myeloid cell-specific deletion of *Hif1a* did not impact on browning marker gene expression in WAT (Fig. 3B). Next, we focused on lipolysis of adipose tissue and performed an *ex vivo* lipolysis assay from eWAT. This assay allowed us to measure the secretion of glycerol from eWAT explants and demonstrated that ASV-B/Hif1a^MC^ mice mobilize fat more efficiently than ASV-B WT mice (Fig. 3C). Finally, we checked whether serum levels of triglycerides were increased. Here, under *ad libitum* food intake conditions, triglyceride levels were found elevated in ASV-B mice compared to controls. However, ASV-B WT and Hif1a^MC^ mice exhibited similar triglyceride levels (Fig. 3D).

**Figure 3.**
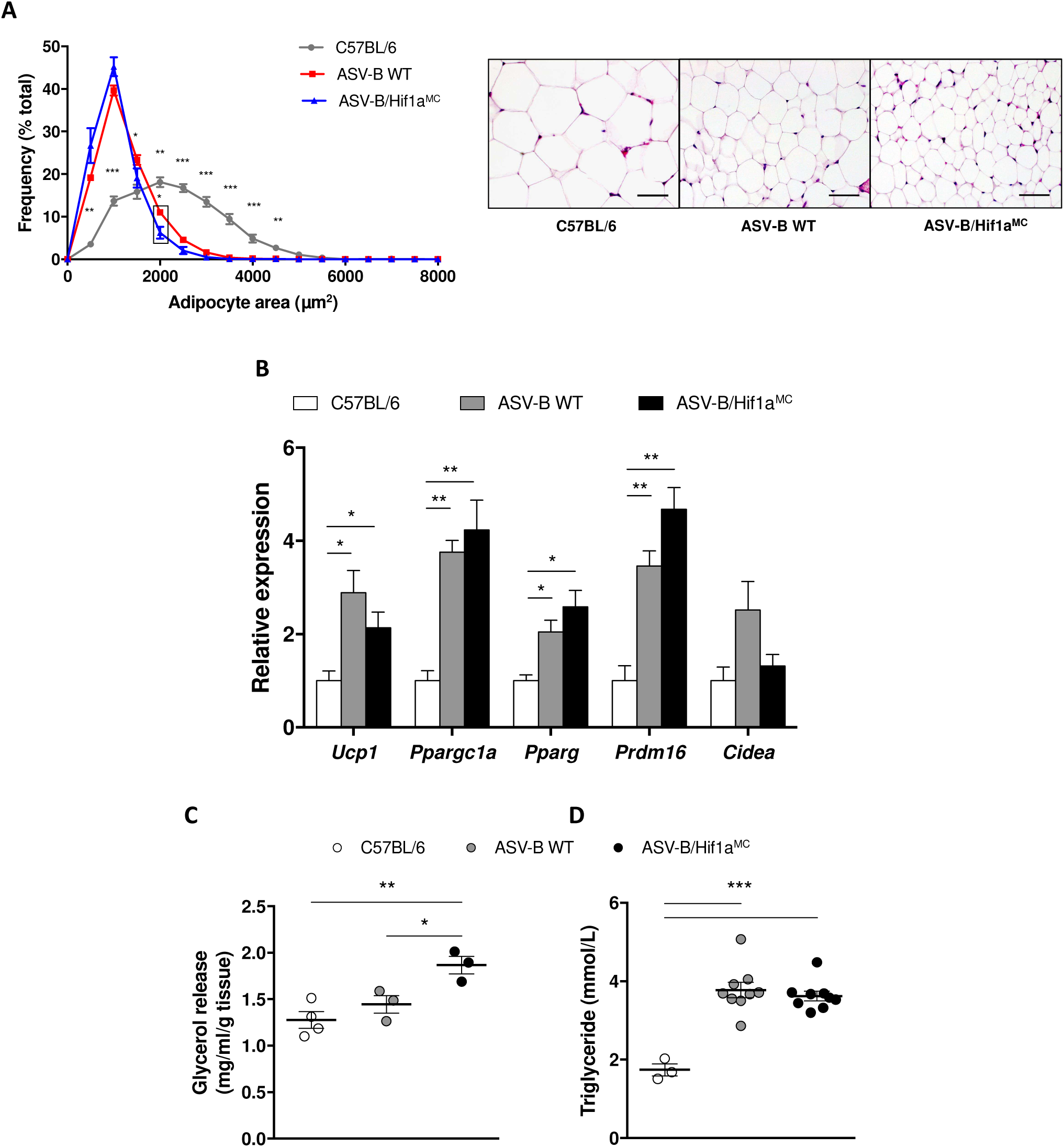
Lipolysis and browning occur in adipose tissue of ASV-B mice. **A**, adipocyte cell size analysis in 16 weeks old C57BL/6 (n = 5), ASV-B WT (n = 4) and Hif1a^MC^ (n = 4) mice. Representative H&E images of eWAT of 16 week old mice (right side); scale bars 50 μm. **B**, mRNA levels of browning marker genes *(Ucp1, Ppargcla, Pparg, Prdm16, Cidea)* as determined by qPCR in eWAT of 16 week old C57BL/6 (n = 3), ASV-B WT (n = 3) and Hif1a^MC^ (n = 4) mice. **C**, glycerol release from eWAT of 16 week old C57BL/6 (n = 4), ASV-B WT (n = 3) and Hif1a^MC^ (n = 3) mice as measured via *ex vivo* lipolysis assay for 2 hours. **D**, triglyceride levels in serum of C57BL/6 (n = 3), ASV-B WT (n = 9) and Hif1a^MC^ (n = 9) mice. Data show means with s.e.m. *P < 0.05; **P < 0.01; *** *P* < 0.001 according to one-way ANOVA followed by Tukey post hoc test.

### Neither tumor load, pro-inflammatory cytokine expression nor hypothalamic activation underlie the enhanced fat loss in ASV-B/Hif1a^MC^ mice

Having confirmed the unexpected aggravation of cancer-associated VAT loss in ASV-B/Hif1a^MC^ mice, we next sought to identify the underlying mechanism(s). One obvious explanation would be an effect of the myeloid cell-specific *Hif1a* deletion on HCC formation, as we have observed for intestinal tumors (24). However, neither μCT-based quantification of liver volume (Fig. 4A), gross liver weight, nor histology-based measurements of tumor load (Fig. 4B) displayed a difference between WT and ASV-B/Hif1a^MC^ mice. As the hypothalamus is able to control lipid uptake and mobilization in WAT (25), we analyzed the activation state of neurons in the nucleus arcuatus (ARC) of the hypothalamus (26). Figure 4C shows the number of c-Fos positive (+) cells in the ARC, reflecting the level of recent neuronal activation. The numbers of c-Fos+ cells in HCC-bearing mice are significantly increased, arguing for an elevated activation of the ARC. As the sympathetic nervous system (SNS) has been shown to mediate the effects of the hypothalamus on WAT (27), we measured catecholamine levels in peripheral fat tissue and found a significant increase in ASV-B compared to tumor-free mice (results for noradrenaline shown in Fig. 4D, adrenaline was not detectable). Again, myeloid cell-specific *Hif1a* deletion remained without a significant effect in these experiments. In adipose tissue, macrophages were suggested as an alternative source of catecholamines (28), although contradictory findings were published in later reports (29). Of note, we were not able to detect noradrenaline or adrenaline in supernatants from bone marrow-derived macrophages from WT and Hif1a^MC^ mice, arguing that macrophages are not a likely source for catecholamines in adipose tissue. Next, we stained eWAT for tyrosine hydroxylase (TH), a marker of sympathetic neurons, the cells that synthesize catecholamines in their axons. Quantitative differences were observed neither between control and ASV-B mice nor between ASV-B WT and Hif1a^MC^ mice (Fig. 4E, F). Finally, we determined serum levels of pro-inflammatory cytokines, which have been shown in numerous studies to be positively associated with cachexia (30, 31). Tumor necrosis factor α (Tnf-α), interleukin 6 (Il-6) and interleukin 1β (Il-1β) are the best studied pro-inflammatory cytokines among these and they can be secreted by macrophages (17). ASV-B WT mice showed significantly increased serum levels of Il-6, Il-1β and Tnf-α compared to tumor-free control mice (Fig. 4G). What’s more, serum levels of Il-1β were significantly reduced while Il-6 and Tnf-α showed a tendency for decrease in ASV-B/Hif1a^MC^ mice. Collectively, these results suggest a functional interplay of pro-inflammatory cytokines and the hypothalamus-peripheral sympathetic nervous system axis in regulating tumor-associated lipolysis in ASV-B mice.

**Figure 4.**
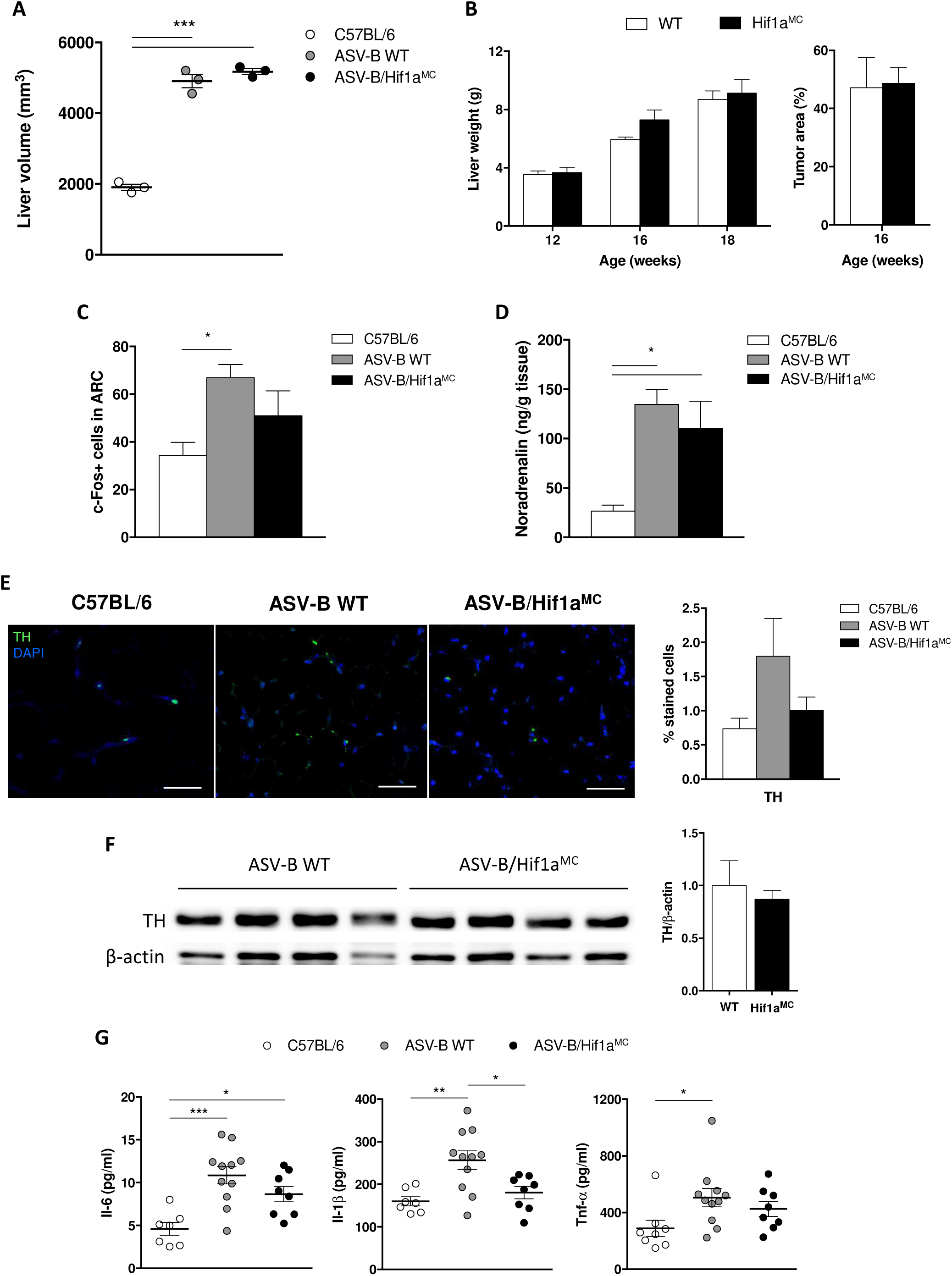
Analysis of potential mechanisms for enhanced fat loss in Hif1a^MC^ mice. **A**, liver volume calculation using μCT images *(n* = 3 per group). **B**, livers were weighed at indicated time points (n = 7, 4, 4 at 12, 16, and 18 weeks, respectively) and tumor area assessment were performed at 16 week old ASV-B WT and Hif1a^MC^ mice (n = 3 per group). **C**, number of cFOS+ cells in arcuate nucleus (ARC) of the hypothalamus of C57BL/6 (n = 3), ASV-B WT (n = 5) and Hif1a^MC^ (n = 3) mice at 16 weeks of age. **D**, noradrenalin levels in eWAT of 16 week old C57BL/6 (n = 3), ASV-B WT (n = 5) and Hif1a^MC^ (n = 4) mice. **E**, (left) representative images of immunofluorescence staining of TH in eWAT from C57BL/6 (n = 5), ASV-B WT (n = 4) and Hif1a^MC^ (n = 5) mice, scale bars 50 μm; (right) quantification of staining, stained cells calculated as relative percentage of all counted cells. **F**, western blot of TH in eWAT from 16 week old ASV-B mice. **G**, serum inflammatory cytokine levels in control (n = 8), ASV-B WT (n = 11) and Hif1a^MC^ (n = 8) mice. Data show means with s.e.m. *P < 0.05; **P < 0.01; *** *P* < 0.001 according to one-way ANOVA followed by Tukey post hoc test (**A**, **B**, **C**, **D**, **E**, **G**) and unpaired Student’s i-test (**B**, **F**).

### Abundance of adipose tissue macrophages is controlled by *Hif1a*

Macrophages have been shown to be important for adipose tissue homeostasis and can be recruited to and accumulate in adipose tissue after lipolysis, where they take part in local lipid regulation (32). Against this background, we characterized different biological aspects of adipose tissue macrophages (ATM) in ASV-B mice. As it was shown that alternatively activated macrophages (AAMs) predominate under conditions of lipid mobilization (33), we decided to analyze macrophage polarization in our model. We applied different experimental approaches, none of which showed a significant effect of *Hif1a* deletion on polarization of adipose tissue macrophages (Fig. 5A, B). Next, we determined ATM abundance via immunohistochemistry against F4/80. Interestingly, while a significant increase in ATM number was noted in ASV-B WT animals, this was completely inhibited upon *Hif1a* deletion (Fig. 5C). Finally, we sought to address a possible contribution of ATM proliferation in our setting. As can be seen in figure 5D, loss of *Hif1a* resulted in a significant decrease of Ki67-positive ATMs, arguing for a functional importance of HIF-1α for ATM proliferation that has not been previously appreciated.

**Figure 5.**
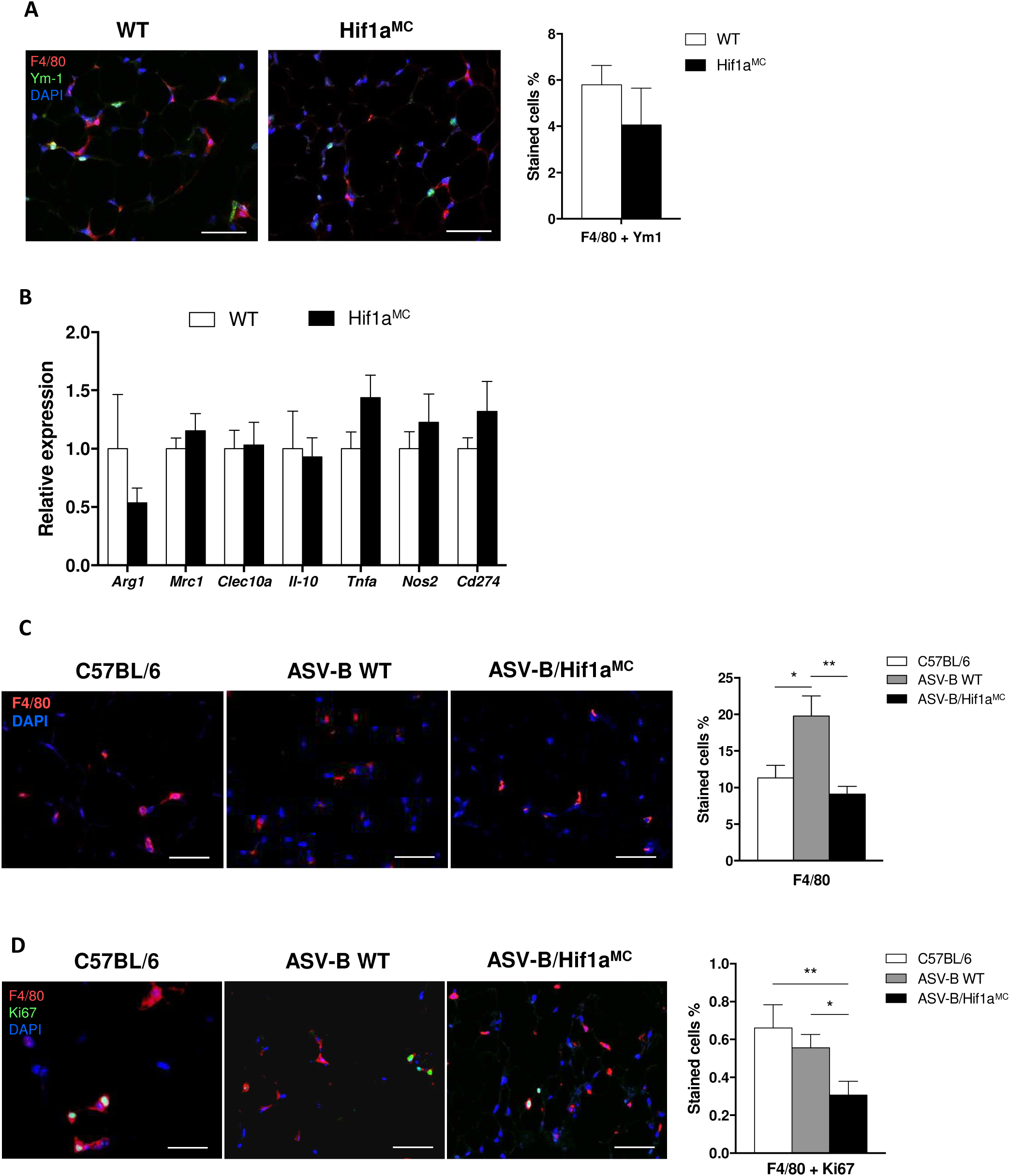
Macrophage phenotype and proliferation in visceral adipose tissue. **A**, (left) representative images of F4/80 and Ym-1 immunofluorescence in eWAT from ASV-B WT (n = 4) and Hif1a^MC^ (n = 5) mice, scale bars 50 μm; (right) quantification of F4/80/Ym-1 double staining, stained cells calculated as relative percentage to all counted cells. **B**, mRNA expression analysis of markers for alternatively and classically activated macrophages in eWAT from 16 week old ASV-B WT (n = 4) and Hif1a^MC^ (n = 5) mice. **C**, (left) representative images of immunofluorescence staining of F4/80 in eWAT from C57BL/6 (n = 5), ASV-B WT (n = 4) and Hif1a^MC^ (n = 5) mice, scale bars 50μm; (right) quantification of staining, positive stained cells calculated as relative percentage of all counted cells. **D**, (left) representative images of immunofluorescence staining of F4/80 and Ki67 in eWAT from C57BL/6 (n = 6), ASV-B WT (n = 4) and Hif1a^MC^ (n = 5) mice, scale bars 50μm; (right) quantification of staining, double positive stained cells calculated as relative percentage of all counted cells. Data show means with s.e.m. *P < 0.05; **P < 0.01 according to unpaired Student’s T-test (**A**, **B**) and one-way ANOVA followed by Tukey post hoc test (**C**, **D**).

### Quantification of visceral adipose tissue in human HCC patients

The robust fat loss of ASV-B mice raised the question about the relevance of this phenotype for the human situation. To this end, we made use of a cohort of HCC patients (n=41, patient characteristics are given in materials and methods) without underlying liver cirrhosis as the ASV-B model is also not associated with hepatic fibrosis (15). Interestingly, 34% of the patients indeed displayed low visceral adipose tissue as determined by the L3 VAT index (Fig. 6). Low L3 VAT index was significantly associated with low body mass index (*P* < 0.001), young age (*P* = 0.017) and female sex (*P* = 0.027), while no association was found with tumor stage.

**Figure 6.**
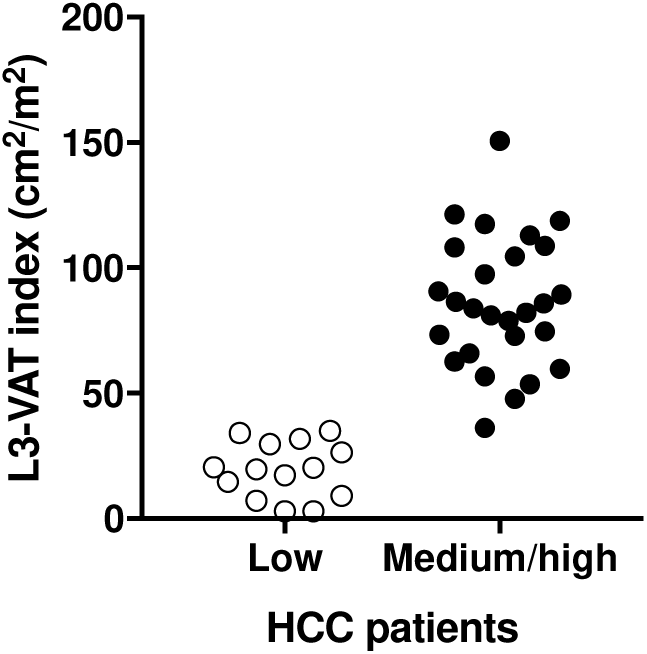
A subgroup of human HCC patients shows a reduced amount of visceral adipose tissue. Distribution of 41 patients according to L3-VAT index which was calculated from CT images of patients. The patient group was divided into tertiles for L3-VAT index, and the lowe tertile was compared to the medium/high tertile. The low tertile L3-VAT index was defined by a cutoff value of 35.

## Discussion

Clinical care of patients with hepatocellular carcinoma is characterized by various challenging obstacles. Co-morbidities of chronic liver disease, limited resectability due to cirrhosis-associated reduction of functional liver reserve and the stout therapy resistance of HCC are among the most widely recognized impediments. For the longest time, the frailty and pronounced muscle weakness that characterizes the majority of HCC patients received much less attention. This has significantly changed in recent years as reports were published from independent groups about the important role of sarcopenia and cachexia in predicting clinical course and response to therapy of HCC patients (4, 34-36). Cachexia is widely considered to be a multi-factorial syndrome with various manifestations throughout the whole body. The causal pathogenesis of cachexia is very complex and involves a plethora of organs, cell types, hormones, cyto-/chemokines, growth factors and interorgan crosstalks (37, 38). To better understand the molecular and cell biological mechanisms that govern cachexia, model systems are needed that recapitulate the syndrome on a whole organism level (39).

Here, we identify a robust cachexia phenotype in the well-established ASV-B mouse HCC model (40). Various characteristic aspects of cachexia were noted in ASV-B mice, e.g. loss of skeletal and heart muscle as well as adipose tissue mass over time, enhanced pro-inflammatory cytokine expression in blood, anemia (Suppl. Fig. 3) and weight loss. While a number of animal models are available to study cancer-associated cachexia (41), only one is widely used with respect to HCC-associated CAC: The rat ascites hepatoma Yoshida AH-130 model (42). This model proved of great value to test the anti-cachexia efficacy of various agents *in vivo*. However, it does not adequately mirror the pathogenesis of HCC as its heterotopic nature does not reflect the liver microenvironment. Furthermore, the AH-130 cells have been established more than 60 years ago (43) and it is reasonable to assume that since then they have acquired a lot of additional changes with unknown relevance for HCC pathogenesis. Recently, it was reported by the group of Erwin Wagner that 50% of mice harboring DEN (di-ethyl-nitrosamine)-induced HCCs developed signs of cachexia at 16-18 months of age (22). While this is a very interesting finding that significantly expands the models available to study HCC-associated CAC, the long time span in combination with a penetrance of 50% are surely obstacles against widespread use of this model. Against this background it is important to note that in ASV-B mice, we were able to detect signs of cachexia with 100% penetrance as early as 12 weeks of age. We therefore consider the ASV-B model to be a powerful addition to the available methodology enabling a better understanding of the mechanistic underpinnings that underlie HCC-associated CAC.

Loss of adipose tissue is a well-known aspect of cancer-associated cachexia and has been noted in animal models as well as samples from patients with various types of cancer (44). It has been reported that adipose tissue loss precedes muscle wasting (45) and that inhibition of the former is able to slow down the latter (14). Reduced peripheral fat content in CAC can be the result of three different processes in adipocytes: Lipid uptake, intracellular *de novo* lipogenesis and lipid release (3). Our finding of visceral adipose tissue wasting in ASV-B mice is well in line with various other murine CAC models (44) and also with the rat hepatoma Yoshida AH-130 model (11). To the best of our knowledge, we are the first to report a functional significance of macrophages for HCC-associated fat loss. Intercrosses of ASV-B with mice showing defective myeloid cell-mediated inflammation resulted in aggravated fat loss. Via immunohistochemistry we could show greater macrophage abundance in AT from wildtype tumor-bearing mice while this phenotype was completely inhibited in KO mice. This led us to hypothesize that HCC-induced AT mobilization results in macrophage influx, ultimately inhibiting lipid release. This would be well in line with earlier reports showing macrophage-mediated suppression of lipid mobilization from AT in response to fasting and pharmacologically induced lipid release in mice (46). Greater macrophage numbers have also been reported in adipose tissue of patients with CAC (47), underscoring the need to better understand the functional significance of these cells for cancer cachexia.

To analyze the role of macrophages for HCC-associated cachexia we made use of a mouse model displaying a defect in myeloid cell-mediated inflammation (19). This is achieved by conditional deletion of *Hif1a* in cells of late myeloid differentiation via the Cre/loxP system (48). *Hif1a* encodes for the transcription factor hypoxia-inducible factor 1α (HIF-1α), the principle mediator of the cellular response to hypoxia (49). HIF-1α target genes control virtually all aspects of the hypoxic response, e.g. erythropoiesis, angiogenesis, glucose metabolism and cell cycle modifications. Inactivation of *Hif1a* in myeloid cells severely impairs energy generation, leading to robustly compromised cellular function and defective myeloid cell-mediated inflammation (19). Our finding of reduced macrophage infiltration into adipose tissue during HCC-associated cachexia is well in line with earlier findings by us and others, demonstrating impaired chemotaxis and migration of *Hif1a*-deficient macrophages, neutrophils and eosinophils in a wide range of underlying pathologies (19, 50). The important question as to the molecular stimuli that attract macrophages to adipose tissue in HCC-bearing mice remains to be addressed in the future. Published work points towards adipocyte-secreted chemokines, e.g. MCP-1/CCL2, and free fatty acids (FFAs) (46, 51). Of note, expression of the MCP-1 receptor CCR2 on monocytes (52) and signal transduction induced by TLR4 (53), the putative cellular receptor for FFAs (54), are strongly influenced by HIF-1α, potentially explaining the reduced macrophage abundance in adipose tissue upon *Hif1a* deletion. In addition to macrophage abundance, our results show reduced proliferation of adipose tissue macrophages in conditional *Hif1a* knock-out mice. While local proliferation of macrophages has been shown in adipose tissue inflammation associated with obesity (55), we are the first to report local macrophage proliferation in the setting of CAC-associated fat loss. Furthermore, a functional role of HIF-1α for macrophage proliferation has thus far only been reported for bovine macrophages after infection with the parasite *Theileria annulata* (56) and not for murine myeloid cells. Admittedly, the percentage of local macrophages that proliferate is rather small. Hence, the functional significance of this observation for CAC-induced lipolysis remains elusive and needs to be validated in future studies.

In line with the mouse model data we were able to show that a subgroup of HCC patients displays a reduced amount of visceral adipose tissue. This strongly suggests that HCC is also capable of inducing fat mobilization from peripheral stores in human patients. To prove this convincingly, one would have to perform longitudinal patient studies, i.e. analyzing the same patient at different stages of disease progression, a venture out of scope of the current project. In other cancer types, e.g. pancreatic adenocarcinoma, adipose tissue loss is a well-known phenomenon with clinical relevance as it is able to predict survival (57). The functional importance of adipose tissue loss for the clinical course of HCC needs to be addressed by future studies with larger patient cohorts. In light of the leading role of adipose tissue loss for the sequence of events characterizing cancer cachexia (14), a better understanding of the mechanisms driving HCC-associated fat mobilization is warranted.

## Materials and Methods

### Animals

Hepatocyte-specific expression of the SV40 large T oncoprotein in ASV-B mice was achieved by the antithrombin III promoter (15). Only male mice develop tumors as the transgene is located on the Y chromosome. ASV-B mice (pure C57BL/6J background) were further crossed with mice with both alleles of *Hif1a* gene flanked by loxP sites at exon 2 (*Hif1a* +^f^/+^f^). Myeloid cell-specific knock-out of *Hif1a* was achieved by breeding ASV-B male *Hif1a* +^f^/+^f^ mice with female *Hif1a* +^f^/+^f^ mice expressing Cre recombinase driven by the lysozyme M promoter. In our study, we used male ASV-B/Hif1a +^f^/+^f^ mice, additionally positive for Cre expression (ASV-B/LysCre+/Hif1α +^f^/+^f^), as knock-outs (named ASV-B Hif1a^MC^) and Cre-negative littermates as wildtype (WT). C57BL/6J male mice were used as controls. All animals were maintained in a specific pathogen-free facility. Mice were given water and standard rodent chow *ad libitum* and were kept at constant room temperature with a 12 hour light/dark cycle. All experiments were approved by local authorities (LAGESO Berlin and LANUV Düsseldorf, Germany) and conducted in accordance with the national and institutional guidelines for care, welfare and treatment for animals.

### Statistical Analysis

The statistical analyzes of data were carried out with Student’s t-test or by One-Way ANOVA, followed by appropriate corrections or post-hoc tests as indicated in the figure legends. Statistical analyzes were carried out with GraphPad Prism 6 software (GraphPad, CA, USA). Data are presented as mean + SEM and the asterisks in the graphs indicate statistically significant changes with *p* values: * *P* < 0.05, ** *P* < 0.01, and *** *P* < 0.001.

**Financial Support:** Research in the Cramer lab was supported by grants from the German Research Foundation (Cr 133/2-1 until 2-4; Cr 133/3-1) and the German Cancer Aid (109160). We gratefully acknowledge support by the European Research Council to Twan Lammers: ERC-StG NeoNano (309495) and ERC-PoC CONQUEST and PIcelles (680882 and 813086). Marco Koch and Henrike Horn receive funding from the German Research Foundation (Collaborative Research Centre 1052 “Obesity Mechanisms”, project CRC1052/A7).

**Conflict of Interest Statement:** The authors declare no potential conflicts of interest.

## Acknowledgements

We are indebted to Dr. Maaike Oosterveer (Department of Pediatrics and Laboratory Medicine, University Medical Center Groningen, The Netherlands) for help with the *ex vivo* lipolysis assay. We are grateful to Dr. Thomas Ritz (University Hospital Aachen) and Prof. Dr. Thomas Longerich (University of Heidelberg) for help with multi-spectral immunofluorescence imaging. We thank Mrs. Constance Hobusch for excellent technical support.

## Supplementary Materials and Methods

### Organ harvest

ASV-B mice were sacrificed at the age of 12, 16, and 18 weeks. 16 week old C57BL/6J male mice were used as control for tissue weights. Blood serum, liver, epididymal white adipose tissue (eWAT), skeletal muscle (gastrocnemius (GC), soleus, tibialis anterior (TA) and extensor digitorum longus (EDL), and heart were collected and weighed after sacrifice.

### Body weight and composition

Body weight, food and water intake were measured weekly. Body composition was analyzed via nuclear magnetic resonance (NMR) spectroscopy device EchoMRI-700™ (Echo Medical Systems, Houston, TX) once a week to measure total body fat and lean mass.

### *In vivo* μCT Imaging

*In vivo* μCT imaging of normal C57BL/6, ASV-B WT and ASV-B Hif1a^MC^ mice was performed using a dual-energy gantry-based flat-panel microcomputed tomography scanner (TomoScope 30s Duo, CT Imaging, Erlangen, Germany). The dual-energy X-ray tubes of the μCT were operated at voltages of 40 and 65 kV with currents of 1.0 and 0.5 mA, respectively. To cover the entire mouse, three sub-scans were performed, each of which acquired 720 projections with 1032 × 1012 pixels during one full rotation with durations of 90 s. Animals were sacrificed just before imaging. After acquisition, volumetric data sets were reconstructed using a modified Feldkamp algorithm with a smooth kernel at an isotropic voxel size of 35 μm. The fat-containing tissue regions, which appear hypo-intense in the μiCT data, were segmented using an automated segmentation method with interactive correction of segmentation errors (1). The volumetric fat percentage was computed as the ratio of (subcutaneous and visceral) fat volume to the entire body volume.

### RNA isolation and qPCR

Total RNA from snap frozen eWAT of 16 weeks old animals was isolated using NucleoZOL (Macherey Nagel, Düren, Germany) and reverse transcription was performed using Maxima Reverse Transcriptase together with Oligo(dT)18 Primers, Random Hexamer Primers and dNTP Mix (Thermo Fisher Scientific, Langerwehe, Germany). Quantitative real-time polymerase chain reaction (qPCR) was performed using Applied Biosystems 7500 Real-Time PCR System in 96-well format, each reaction containing 15 ng cDNA, 0.3μM specific primer and 1x *Power* SYBR Green Master Mix reagent (Applied Biosystems, Bleiswijk, The Netherlands). Primers for *Ucp1, Pgc1a, Pparg, Prdm16, Cidea* and *Mrc1* were used as described before (2, 3) as well as the following primers: *B2m* F: 5’-TTCTGGTGCTTGTCTCACTGA-3’, R: 5’-CAGTATGTTCGGCTTCCCATTC-3’; *Arg1* F: 5’-CTCCAAGCCAAAGTCCTTAGAG-3’, R: 5’-AGGAGCTGTCATTAGGGACATC-3’; *Clec10a* F: 5’-GGCACAAAACCCAGCAAGAC-3’, R: 5’-TGGGACCAAGGAGAGTGCTA-3’; *Il10* F: 5’-GCTCTTACTGACTGGCATGAG-3’, R: 5’-CGCAGCTCTAGGAGCATGTG-3’; *Tnfa* F: 5’-CCATTCCTGAGTTCTGCAAAGG-3’, R: 5’-AGGTAGGAAGGCCTGAGATCTTATC-3’; *Azgp1* F: 5’-ACACTACAGGGTCTCACACCT-3’, R: 5’-TCGCTGCACGTAGACCTTTT-3’; *Lipe* F: 5’-TGTCACGCTACACAAAGGCT-3’, R: 5’-GGTCACACTGAGGCCTGTC-3’. Relative mRNA expressions were calculated using the comparative delta-CT method and normalized to *B2m*.

### Cytokine measurement

Blood was collected from sacrificed mice via inferior vena cava using a 22 G needle and transferred to serum-gel Z tubes (Sarstedt, Germany), allowed to clot for 30 min at room temperature and centrifuged at 10.000g for 5 min. The serum was collected and stored frozen until use. To detect interleukin-1 beta (Il-1β), interleukin-6 (Il-6), and tumor necrosis factor-alpha (Tnf-α) simultaneously, Bio-Plex Pro™ mouse cytokine Th17 panel A 6-Plex kit (Bio-Rad, Germany) was used according to manufacturer’s instructions. Samples were diluted at 1:2 and the fluorescence measurement of the beads was done with the Qiagen LiquiChip 200 workstation (Hilden, Germany). Cytokine concentrations were calculated using Bio-Plex Manager software (Bio-Rad, Hercules; CA USA).

### Immunohistochemistry and tissue analysis

Mice were sacrificed and tissues were fixed in 10% formalin overnight, followed by dehydration and embedding in paraffin. For histopathological evaluation, two-micrometer-thick eWAT or liver sections were stained with hematoxylin and eosin (H&E). For adipose tissue, pictures of representative areas from each section in 200x magnification were taken and Adiposoft software was used to calculate cell size of 35 images per group in total. Minimal 20 μm and maximum 100 μm thresholds were set for automated measurement of adipocyte diameter followed by manual correction. A frequency distribution was calculated for each group. Total adipocyte number within the distribution was subsequently calculated and the frequency was converted to a percentage of total adipocytes counted. For analyses of tumor areas, H&E stained sections of ASV-B livers were used. Two tissue sections per mouse were used for evaluation. Images were taken using Axiocam 506 mono (Carl Zeiss) and tumor areas were quantified by ImageJ.

### Immunohistochemistry of murine hypothalamus

Free floating coronal brain sections of 40 μm thickness were cut on a microtome (Leica VT1200) and stored in 0.02 M PBS + 0.09 % sodium azide until staining. For c-fos immunolabeling we used a modified version of a published protocol (4). Slices containing the arcuate nucleus (ARC) were blocked and permeabilized in 0.02 M PBS + 0.3 % Triton X-100 + 5 % normal goat serum (NGS, Jackson ImmunoResearch) for 60 minutes. The slices were incubated in rabbit polyclonal anti-cfos-antibody (1:10.000, Synaptic Systems, #226003) in PBS + 0.3 % Triton X-100 + 3 % NGS for 5 nights at 4°C. After three washes in PBS + 0.3 % Triton, brains were incubated with goat secondary antibodies raised against rabbit, conjugated to Alexa fluor 488 (1:500, Invitrogen, #A11034) in PBS + 0.3 % Triton X-100 + 3 % NGS for 60-90 minutes at room temperature. To visualize nuclei, 4',6-Diamidin-2-phenylindol (DAPI, 1:10.000) was added for 5 minutes to one of the final three washing steps in PBS. Sections were mounted on glass slides (Menzel Gläser), embedded in fluorescence mounting medium (Dako) and covered with glass cover slips (Menzel Gläser). Brain sections were imaged using confocal laser scanning microscopy (Zeiss, LSM700), operated by ZEN 2011 SP3 (Zeiss). Images were taken using a 20x Plan-Neofluar objective (NA 0.5) using the same imaging parameters for all images. Optical sections (thickness: 9.8 μm in 488 channel, 9.9 μm in DAPI channel) were acquired in 5 μm steps from each hemisphere of the ARC. All image analysis was carried out in ImageJ (https://imagej.net) using custom macros. One optical section located in the middle of each brain slice was extracted and brightness and contrast were adjusted with the same parameters for all images to improve visibility. Putative c-fos positive cells were counted manually in the Alexa 488 channel using the Cell Counter plugin (https://imagej.net/CellCounter) by an observer blind to the experimental groups. Data from two staining experiments were taken together and least four ARC hemispheres from 2-3 brain sections were analyzed per animal.

### Triglyceride measurement

Sera samples were collected from sacrificed animals as described for cytokine measurement. Triglycerides were measured in the central biochemistry laboratory of the Institute for Laboratory Animal Science, RWTH Aachen University Hospital.

### *Ex vivo* lipolysis

Gonadal fat pads were excised from mice, cut into 20 mg pieces and incubated at 37 °C in Krebs-Ringer medium containing 2% of free-fatty acid BSA. Released glycerol was measured from supernatants after 4 hours incubation using the Glycerol Colorimetric Assay Kit (Cayman) and tissue weights were used for normalization.

### Isolation and stimulation of bone marrow-derived macrophages (BMDM)

Bone marrow was collected from tibiae and femurs of 8-11 weeks old WT and Hif1a^MC^ mice. Red blood cells were lysed with ACK buffer in flushed marrows and cells were seeded on cell culture plates in RPMI supplemented with 10% FBS, 100 U/ml penicillin, 100 μg/ml streptomycin. After overnight incubation, non-attached cells were collected and cultured in RPMI supplemented with 20% FBS and 30% L929-conditioned medium for one week. Differentiated BMDMs were stimulated for 48h with LPS (100 ng/ml, Sigma Aldrich) and IFN-Y (20 ng/ml) for classical activation and with IL-4 (20 ng/ml, both from eBioscience) for alternative activation of macrophages. Media were collected from polarized macrophages and used for catecholamine measurement.

### Catecholamine measurement

Catecholamine amounts were measured with high performance liquid chromatography (HPLC). Snap frozen eWAT samples were thawed and sonicated in 0.3 M perchloric acid for 30 s on ice (200μl/0.1g tissue). Samples were centrifuged at 9.000 rpm for 10 min at 1°C. Supernatants, cleaned from residue, were collected and used for HPLC measurements. Cell culture media collected from BMDMs were directly injected into the system. All measurements were performed by a service laboratory with special expertise in HPLC (Laboratory for Stress Monitoring, Hardegsen, Germany).

### Western Blotting

50 mg eWAT samples were homogenized in 100 μl RIPA buffer containing 10 mM Tris-HCl (pH 7.5), 150 mM NaCl, 0.25% SDS, 1% sodium deoxycholate 1% NP40, 2 mM PMSF, 1 mM DTT; 10 mM NaF, 1 mM Na_3_O_4_ and 2μM Leupeptin and 4.4x10^-4^ TIU/ mg Aprotinin. After sonication, the homogenates were centrifuged for 15 min at 12.000 g, 4°C and supernatants were collected. Total protein content was measured by Lowry assay (DC Protein Assay, Bio-Rad). 40μg protein were separated via SDS-PAGE and transferred to a nitrocellulose membrane. The membrane was incubated overnight with TH antibody (Millipore, AB152) in 5% milk-0.05% TBS-Tween 20 (1: 1.000), and β-actin antibody (Sigma, A5441) (1:5.000) was used as a loading control. Membranes were developed using enhanced chemiluminescence reagent (Perkin-Elmer™, Life Sciences) and visualized by ChemoCam Imager (INTAS, Göttingen, Germany).

### Immunofluorescence staining and quantification

Paraffin sections from eWAT with a thickness of 2 μm were deparaffinized and rehydrated according to standard protocols. Sections were re-fixed by 10 min incubation in 3.5% formalin and antigen retrieval was done by 15 min of cooking at 110°C in Dako target retrieval solution in the Decloaking Chamber (Biocare Medical, Berlin, Germany). After 10 min of blocking in Dako antibody diluent, sections were incubated with F4/80 (eBioscience #14-4801) antibody in a 1:2.000 dilution for 30 min followed by ImmPRESS anti-rat IgG (Vector) incubation for 20 min. For permanent labeling of F4/80 with a fluorescent tag, sections were treated with Opal 570 (Opal 4-color IHC kit, PerkinElmer, 1:50 in amplification reagent) for 10 min. Subsequently, the antibodies were detached from sections by microwave cooking in AR6 buffer (Opal 4-color IHC kit, PerkinElmer), whereas the Opal fluorophore remained fixed to the tissue. For staining of a second marker, sections were directly processed further and the described procedure was repeated. Briefly, sections were blocked again and incubated either with anti-TH (Millipore #AB152, 1:5.000), anti-Ki-67 (Cell Signaling #12202, 1:3.000) or anti-Ym-1 (Stemcell Tech. 60130, 1:5.000). Following anti-rabbit IgG treatment, antigens were labeled with Opal 520 (Opal 4-color IHC kit, PerkinElmer, 1:100 in amplification reagent). After microwave treatment, nuclei were stained with spectral DAPI (PerkinElmer) for 5 min. Tissue sections were covered with Vectashield HardSet antifade mounting medium (Vector Laboratories). Fluorescent signals were detected, separated and recorded using the Vectra 3.0 multiplex imaging system (PerkinElmer). Quantification of signals were performed via inForm automated image analysis software (PerkinElmer).

### L3-Visceral adipose tissue index analysis of HCC patients

Routine CT scans (performed maximally 6 weeks before surgery) from 63 HCC patients of the Department of General, Transplantation, and Visceral Surgery at the University Hospital RWTH Aachen were scheduled for body composition analysis following ethics approval of the local authorities. Abdominal scans were analyzed in a blinded approach in an anonymized format. At first, for each patient, a single slice of CT scan at the level of the third lumbar vertebra (L3) was selected. CT scans with low quality image or missing parts of muscle tissue on the ventral, dorsal, or both lateral edges of the scan were excluded. After L3 selection, 41 patients became eligible for analysis. L3 cross-sectional images were analyzed using sliceOmatic software (version 5.0, TomoVision, Montreal, Canada) to determine adipose tissue area. Tissue Hounsfield unit (HU) thresholds were set up as recommended by the software, -150 to -50 for visceral adipose tissue (VAT) cross-sectional areas (cm^2^) were determined as assigning a specific tag on different tissues. Total area of VAT was normalized for height of the patients to calculate L3-VAT index in cm^2^/m^2^ which provides a powerful estimation of total body VAT mass. Sex-specific cutoff values were established by dividing cohort into tertiles. By the use of the lowest tertile as the reference group in comparison to the middle and high tertiles, statistical analyses were performed.

**Supplementary Figure 1.**
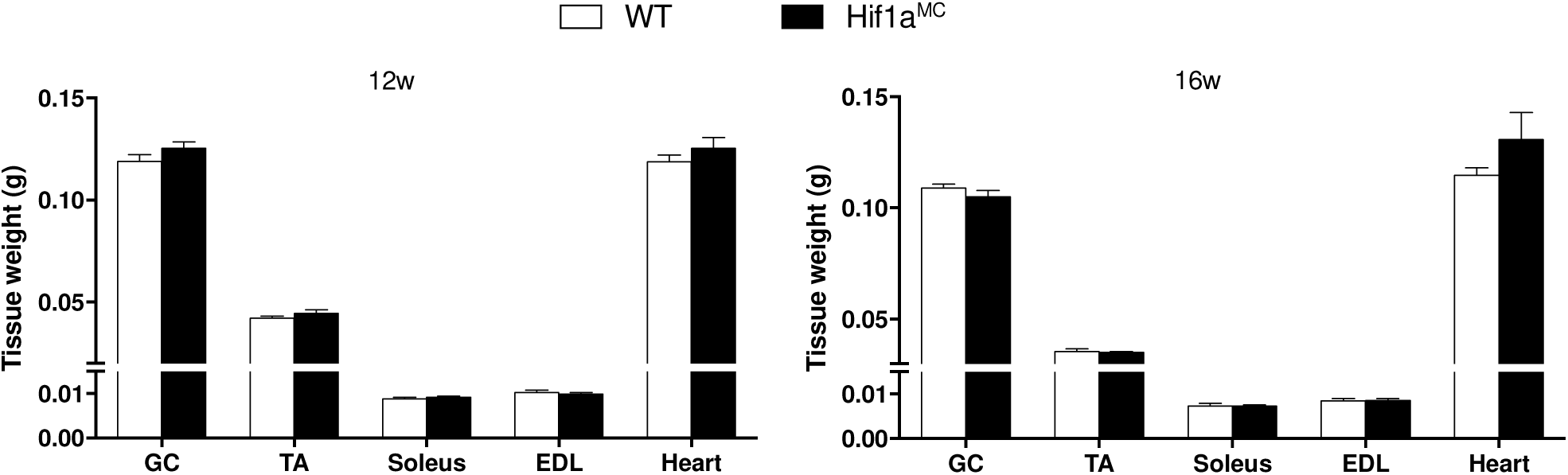
Measurements of skeletal muscle (GC, TA, EDL, soleus) and heart muscle were performed at 12 (right, 12w) and 16 (left, 16w) weeks of age. Data show means with s.e.m. and were analyzed by unpaired Student’s t-test.

**Supplementary Figure 2.**
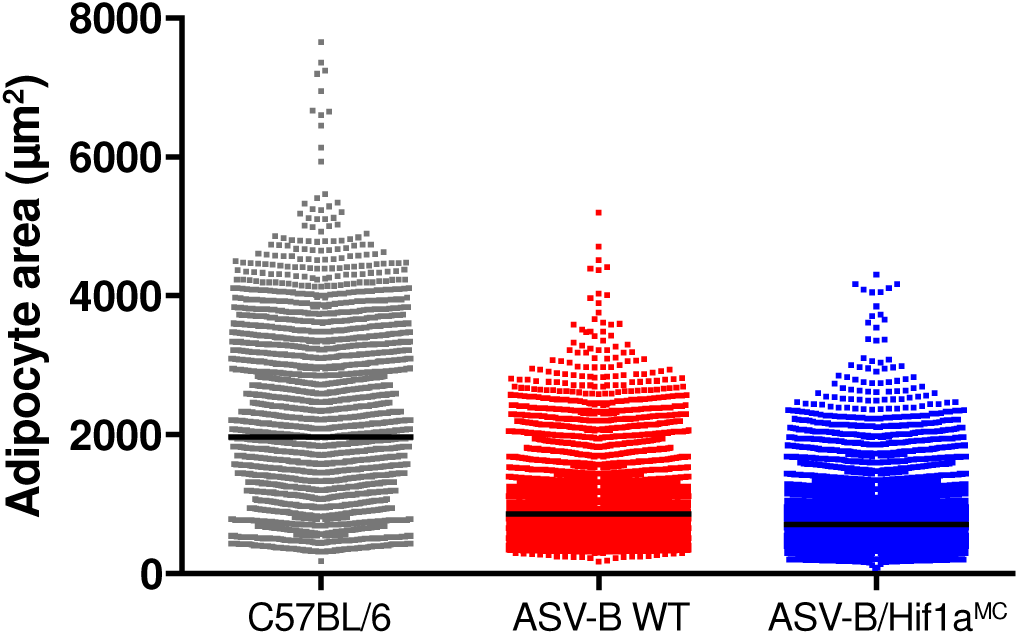
Adipocyte cell area analysis was performed using Adiposoft software as described in Figure 3A and materials and methods. This scatter plot represents the distribution of all adipocytes according to their size together with the mean of each group.

**Supplementary Figure 3.**
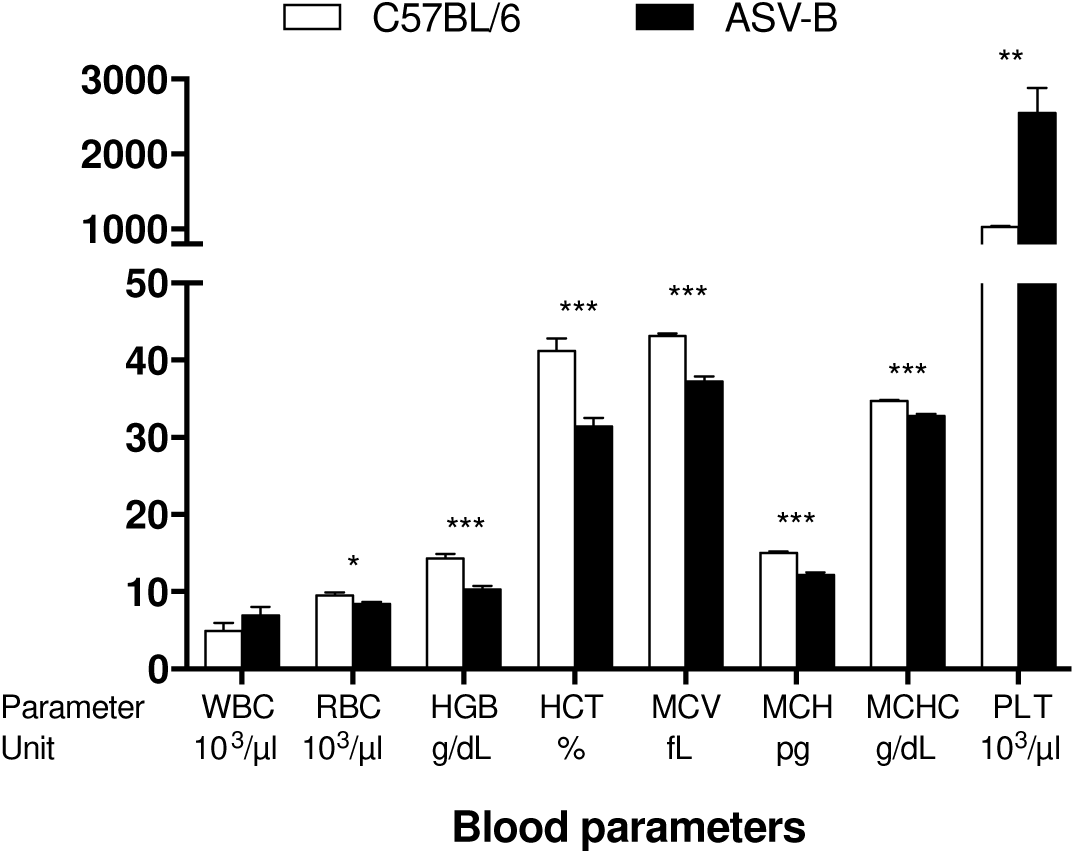
Hematology blood work of ASV-B mice. Complete blood counts in C57BL/6 (n = 5) and ASV-B (n = 6) mice at 16 weeks of age were performed in automated analyzer. WBC: White blood cells, RBC: Red blood cells, HGB: Hemoglobin, HCT: Hematocrit, MCV: Mean corpuscular volume, MCH: Mean corpuscular hemoglobin, MCHC: Mean corpuscular hemoglobin concentration, PLT: Platelets. Data represent means with s.e.m. analyzed by unpaired Student’s *t*-test; \**P* < 0.05; \*\**P* < 0.01; *** *P* < 0.001.

